# Predicting neoadjuvant chemotherapy benefit using deep learning from stromal histology in breast cancer

**DOI:** 10.1101/2022.06.19.496741

**Authors:** Fengling Li, Yongquan Yang, Yani Wei, Yuanyuan Zhao, Jing Fu, Xiuli Xiao, Zhongxi Zheng, Hong Bu

## Abstract

Neoadjuvant chemotherapy (NAC) is a standard treatment option for locally advanced breast cancer. However, not all patients benefit from NAC; some even get worse outcomes after therapy. Hence, predictors for treatment benefit are crucial for guiding clinical decision-making. Here, we investigated the predictive potentials of breast cancer stromal histology via a deep learning (DL)-based approach and proposed the tumor-associated stroma score (TS-score) for predicting pathological complete response (pCR) to NAC with a multi-center dataset. The TS-score is demonstrated to to be an independent predictor of pCR as it not only outperformed the baseline variables and stromal tumor-infiltrating lymphocytes (sTILs) but also significantly improved the prediction performance of the baseline variable-based model. Further, we discovered that unlike lymphocyte, collagen and fibroblasts in stroma were likely associated with poor response to NAC. The TS-score has potentials to be a candidate for better stratification of breast cancer patients in NAC settings.

## Introduction

Neoadjuvant chemotherapy (NAC) is a standard treatment option for patients with locally advanced breast cancer and some large operable tumors [1, 2]. NAC has been shown in clinical trials to reduce the tumor burden and promote the breast-conserving surgery, with patients who achieved a pathological complete response (pCR) having a better prognosis [3]. The pathological response rate, on the other hand, varies among patients who receive this treatment modality, and is primarily determined by their molecular subtypes [4, 5]. The heterogeneity of breast cancer in response to NAC has sparked renewed interests in predictive biomarkers, since these facilitate clinical decision-making at the early stage.

Histological images contain a wealth of tumor phenotypic information and reflect the underlying molecular processes and disease progression, which can provide intrinsic information of diseases for clinic. Subjective evaluation of pathological slides by well-trained pathologists is the gold standard for disease diagnosis and classification. However, pathological diagnosis mainly relies on visible morphological features while the abundance of clinically relevant hidden information that is currently not fully exploited yet. For instance, the Nottingham grading system provides prognostic and predictive information about the breast cancer through pathologist assessment of histological features including nuclear atypia, glandular differentiation, and mitotic count, but manual assessment can be subjective and less reproducible and relies only on limited visible visual features. In recent studies, digital pathology and artificial intelligence (AI) techniques, which enable the extraction of hidden and quantitative information directly from histological images, have showed the potentials of providing clinically useful indicators [6, 7]. Particularly, the introduction of the convolutional neural network (CNN) revolutionized the field of image analysis. Neural networks can distinguish objects by learning features from the training data and effectively solve complex visual tasks [8]. Previous studies on digital pathology have used AI-based image analysis methods for tumor detection [9], tumor grading [10, 11], immunohistochemistry (IHC) scoring [12] and other medical classification tasks [13-15], showing great potentials in clinical application. More recently, deep learning (DL) methods based on medical images were used to develop novel biomarkers that were found to be predictive of the prognosis and chemotherapy response [16-20].

In a previous study, we proposed an image-derived biomarker for predicting pCR in breast cancer, which revealed hidden predictive information from tumor epithelium [20]. Nevertheless, the tumor-associated stroma, also known as the tumor microenvironment (TME), offers numerous potentials for the discovery of novel biomarkers for predicting the disease outcome. Tumor-associated stroma constitutes a suitable microenvironment for tumor growth, progress, and metastasis; the stromal phenotypic information presented on histology reflects the aggregate effect of underlying tumor biological alterations [21]. However, the high heterogeneity and complexity of TME has hampered the research progress on stromal derived biomarkers from histological images. With the employment of AI techniques, several studies have found that the stromal morphological features are predictive of prognosis in breast cancer [18], prostate cancer [19], and colorectal cancer [17]; particularly, Beck et al. proposed that the quantitative information extracted from stroma was fairly predictive for prognosis in breast cancer [18]. Nevertheless, few studies have investigated the potential value of stroma to predict the treatment response to chemotherapy. Although some stromal parameters from manual evaluation, like the tumor-infiltrating lymphocytes and tumor-stroma ratio, have showed some predictive ability to pCR [22-24], abundant hidden information of the stromal morphology still remains to be exploited.

In this study, we aimed to fill this gap by exploring the potential ability of tumor-associated stroma using AI techniques. We hypothesized that a stromal derived biomarker could improve the prediction of pCR in breast cancer. We used DL-based methods to propose a stromal derived biomarker from hematoxylin and eosin (HE) -stained histological images of breast cancer biopsies and evaluated the predictive power in four independent, multi-center datasets.

## Materials and Methods

### Study design

Based on a multi-center study of 1035 breast cancer patients from four independent Chinese hospitals, a new biomarker, called tumor-associated stroma score (TS-score) directly derived from tumor stromal compartment was proposed to predict the treatment response to NAC in patients with primary invasive breast cancer. Histopathological assessment of the resected breast specimens after surgery was chose as the reference standard, and TS-score was compared with baseline clinic-pathological (CP) variables and manual-evaluated TILs derived from tumor stroma. The predictive incremental value of TS-score for predicting pCR was evaluated as well using CP variable-based model as the reference baseline. Besides, we explored the potentially histological pattern of the breast cancer stroma that the TS-score characterized. Our study has been approved by the ethics committee of each participating hospital, and abided with the Declaration of Helsinki before using tissue samples for scientific researches purpose only. The requirements to obtain informed consent from the participants were waived by the ethics committee.

### Patients

The inclusion criteria were as follows: 1) patients with primary invasive ductal breast cancer; 2) patients without distant metastasis; 3) patients receiving four, six, or eight cycles anthracycline and/or taxane-based NAC regimens, and patients with human epidermal growth factor receptor 2 positive diseases (HER2+) underwent targeted HER2 therapy (detail NAC regimens were available in Table S1); 4) patients who have undergone surgical treatment after NAC. On the other hand, patients with HE-stained slides of poor quality including the tissue-processing artifacts like bubbles, discoloration and soiling caused by long storage time, and low tissue volume were removed from our study. Totally, 1035 eligible patients were enrolled, and detailed recruitment flow was shown in Figure 1. The dataset with the largest population was assigned as the primary cohort (PC) for developing the image-derived predictor, and the other three cohorts were used as validation cohorts (V1∼V3).

**Figure 1.**
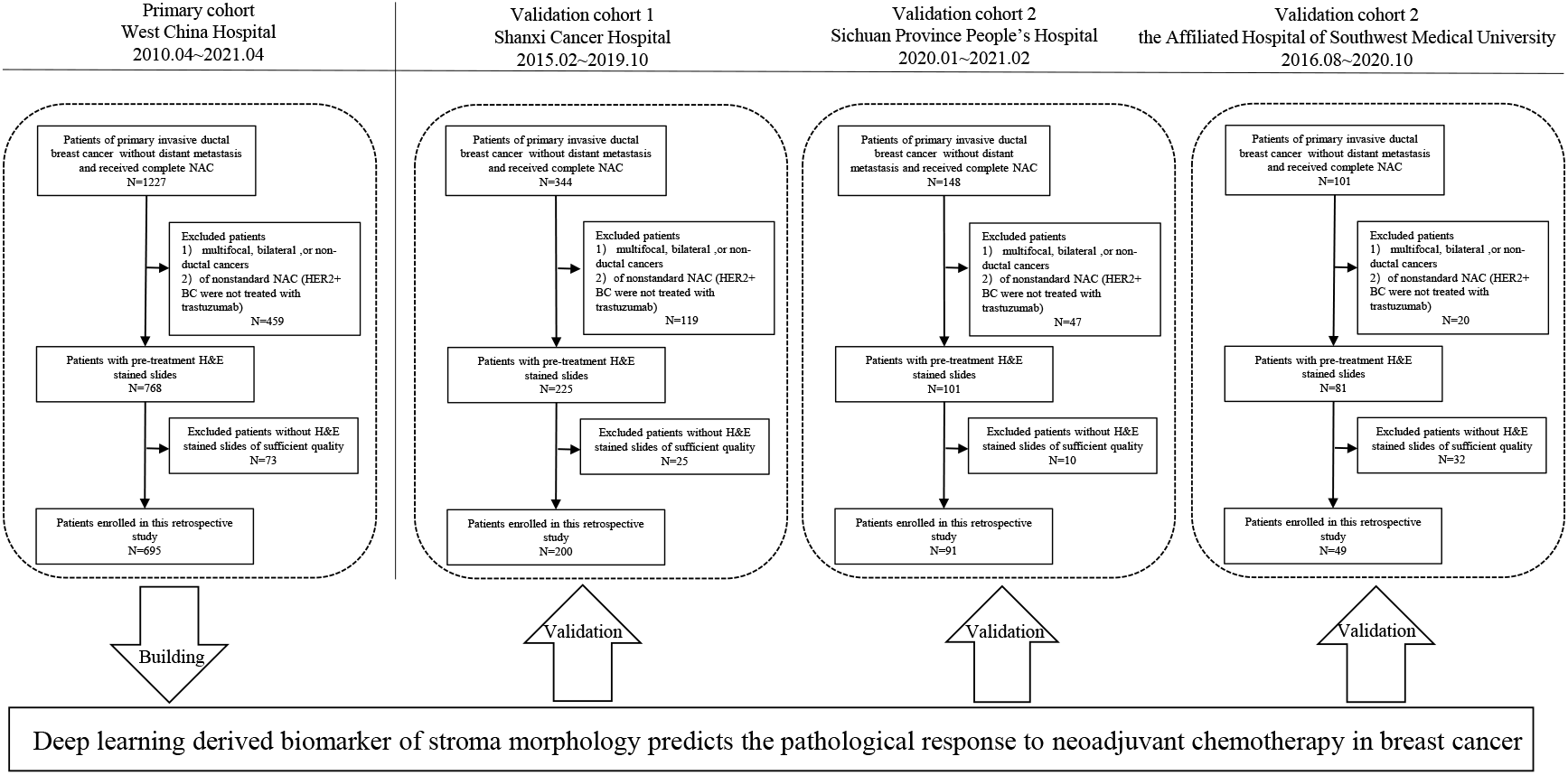
Patients recruitment and study design. 1035 patients out of 1820 with pretreatment H&E stained slides from four Chinese hospitals were included in this study for stroma-derived biomarker development and validation.

### Data and image acquisition

Histological sections and clinic-pathological data were obtained from the corresponding hospitals and delivered to the center laboratory for a unified process. Sections of HE-stained, formalin-fixed, paraffin-embedded breast cancer biopsies were manually reviewed to exclude cases with tissue-processing artifacts or poor staining. Eligible sections were digitally scanned at 40× magnification using a Hamamatsu scanner. Clinical variables such as diagnostic age, tumor size, and clinical lymph node status were gathered from the medical records at each institution, and pathological indicators including ER, PR, HER2, Ki67 results were collected from the pathological diagnostic reports. No less than 1% positive cells for ER/PR IHC examination was defined as ER/PR positive, and ER and/or PR positive breast cancer was classified into hormone receptor positive (HR+) disease. For HER2 status, IHC 3+ and/or amplified by fluorescence *in situ* hybridization (FISH) were regarded as positive, otherwise IHC 0/1+ and IHC 2+ with no amplification by FISH was HER2 negative (HER2-) disease. According to the reference [25], 20% was set as the cutoff point of Ki67, which grouped the cases into the high expression cohort and the low expression cohort. The pathological response to NAC was reviewed at the center laboratory, and patients were classified into pCR group and non-pCR group at each hospital. Here pathological complete response was referred the invasive tumor cells were eliminated at the primary breast site (ypT0).

### Pathological evaluation

Stromal tumor-infiltrating lymphocyte was assessed following the international recommendation guidelines [26]. Briefly, all stromal mononuclear cells within the tumor border, including lymphocytes and plasma cells but not the macrophages and neutrophils, were counted and the percentage of stromal TILs was estimated as a semi-quantitative continuous parameter indicating the density of sTILs. Besides, sTILs was categorized as three grades: low (⩽10%), moderate (11%∼39%), high (⩾40%) [26]. Nuclear grade was assessed based on the Nottingham grading system. Additionally, stromal type classification was performed on patch-level and WSI-level by two well-trained observers following the criteria in previous studies [27]. According to the main component of the stroma, patches/WSIs were classified into collagen dominant (C type), fibroblast dominant (F type), and lymphocyte dominant (L type); cases which did not fall into one of the three categories were as unclassified type. The sTILs, nuclear grade, and stromal type were evaluated on the digital images by two independent observers at the center laboratory, and inconsistent cases were reviewed to reach a consensus.

### Image processing pipeline

We developed a customized image processing pipeline consisting of three main steps: annotation for the region of interest (ROI), training and employment of the epithelium-stroma classifier (E-S classifier, CNN I), TS-score development (Figure 2). Representative tumor regions containing tumor stroma were manually annotated on each WSI, ensuring stroma inside the ROIs was near to the tumor and surrounded by tumor cells (detail illustrations for ROI annotation are available in Supplementary materials) [19, 28, 29]. Images from ROIs were preprocessed and cropped into 233×233 μm square (256 × 256 pixels at 10 × magnification), called “tile”.

**Figure 2.**
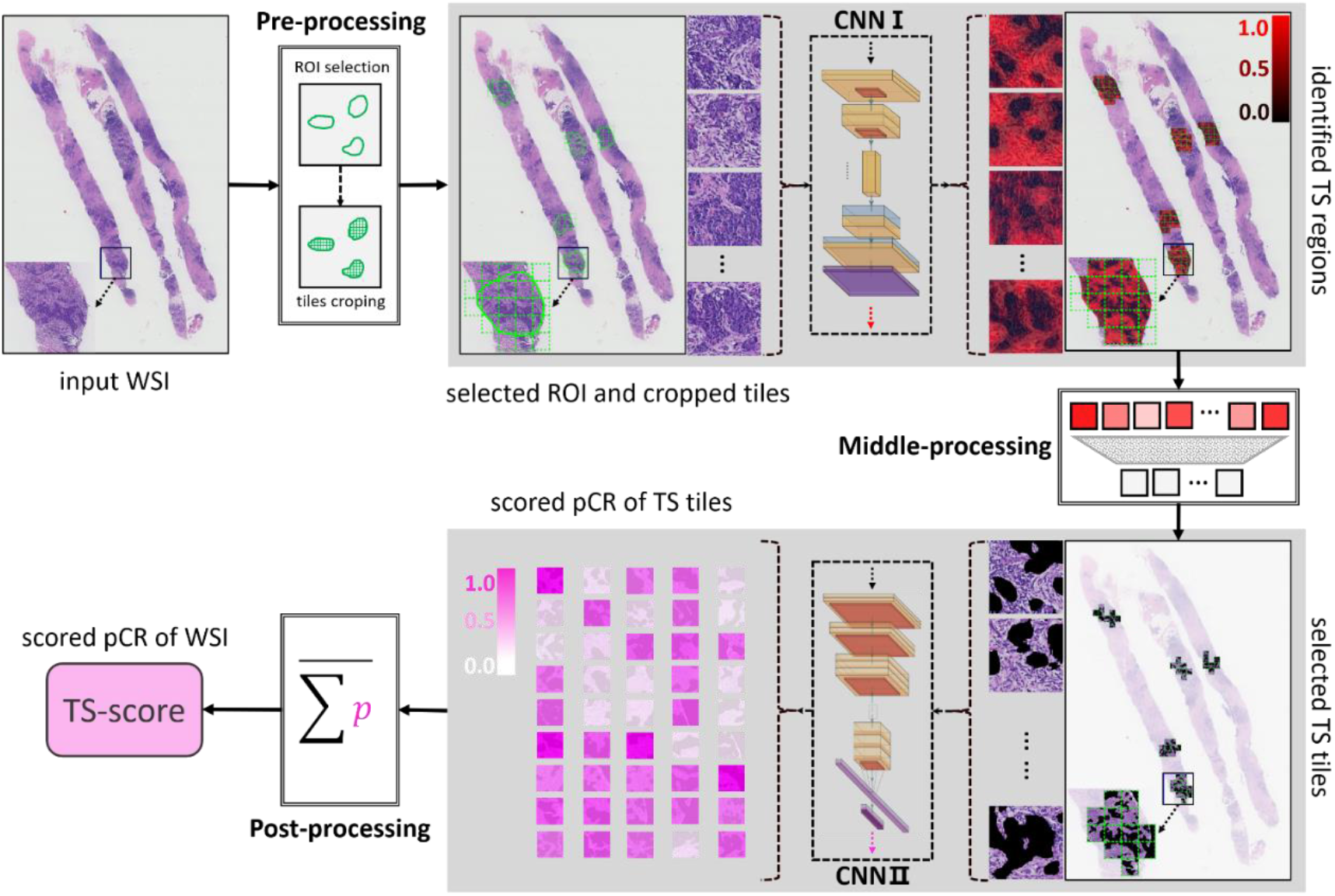
Image processing pipeline to develop a stromal-derived biomarker for predicting pCR. In the pre-processing step, the digitized HE-stained slides were manually annotated and the ROIs were cropped as tiles (256 × 256 pixels at 10 × magnification). With processed by the CNN I (also referring the E-S classifier), stromal pixels within the ROIs were detected and highlighted in red, and the color, red to black, indicates the probability of stroma from high to low. In the middle-processing step, a well-trained observer reviewed all tiles and removed the stromal tiles that did not exactly matched with the ground truth. Stroma tiles with identified by both the CNN I and the human observer were delivered to the CNN II and each tile was assigned with a score indicating the risk of achieving a pCR. In the post-processing step, all tile-level scores of each WSI were summed and the mean value was calculated and named TS-score, which was deemed as a DL-based biomarker derived from the stromal compartment and reflected the risk of pCR for breast cancer.

To concentrate on the tumor stromal compartment, we trained an E-S classifier (CNN I) identifying the stroma from the ROIs (see Supplementary methods). Two annotation strategies for tumor epithelium were used by a well-trained pathological observer to better train the model while using less manual efforts [5], and data with noisy labels (annotations) were fed to train an image semantic segmentation model for the task of identifying the tumor stroma. During the inference by the CNN I, areas inside the ROIs were segmented as epithelial compartment or stromal compartment (see Figure 2 and Figure S2). To avoid the misclassification of stromal regions by the E-S classifier, a well-trained human observer reviewed all regions predicted as stroma and removed tiles of inaccurate classification (see Supplementary Materials and Figure S4). Finally, the stromal areas identified by both the E-S classifier and the human observer were used to develop a stroma-derived signature for predicting the treatment outcome in breast cancer.

### Training and testing of neural networks

To train an E-S classifier, we employed one-step abductive multi-target learning with diverse noisy samples (OSAMTL-DNS) to learn the labeled noise samples more effectively [30], as shown in Figure S2. More details about training and employment of CNN I were available in Supplementary Materials. The tiles with exact stromal mask from CNN I were delivered the CNN II to perform TS-based scoring; each tile inherited the label (pCR vs. non-pCR) from the corresponding WSI, that was, patient-level. To develop a stroma-based biomarker for predicting pCR, the Inception-V4 was selected as the base DL architecture [31]. Weighted cross-entropy [32] and stochastic gradient descent (SGD) [33] were used in optimization. Moreover, we used the fast ensemble DL strategy to further boost the optimized of the prediction part of CNN II [34-36]. After scoring all tiles, an averaged value from all the tiles of each WSI was calculated as the TS-score, which reflected the risk of obtaining a pCR for an individual patient (Figure 2 and Figure S5).

### Statistical Analyses

Comparisons among cohorts and between pCR/non-pCR groups were made with the Pearson χ^2^ test or Fisher’s test for qualitative variables (Table 1, Table S3), while the t test or Mann–Whitney U test was used for continuous factors (Figure 5). Univariate and multivariate logistic regression methods were used to investigate the correlations between factors and pCR in PC and VCs. AUCs and 95% confidence interval (95%CI) were used for evaluating the prediction performance, and the Delong test was used to compare the difference between AUCs [37]. The AUC of bootstrap analysis (100 repetitions) was performed to estimate the CI in the validations while the 5-fold cross validation was used in the primary cohort. All the statistical analysis was two-sided and P-value is less than 0.05 indicating statistical significance. The statistical analyses were performed using SPSS software, version 25.

**Table 1.** Demographic and clinic-pathological characteristics.

|  | PC<br>(n=695) | V1<br>(n=200) | V2<br>(n=91) | V3<br>(n=49) | P |
| --- | --- | --- | --- | --- | --- |
| **Age at diagnosis** |  |  |  |  | **<0.001** |
| <50 | 402 (57.8) | 71 (35.5) | 37 (40.7) | 26 (53.1) |  |
| ≥50 | 293 (42.2) | 129 (64.5) | 54 (59.3) | 23 (46.9) |  |
| **cT (%)** |  |  |  |  | **<0.001** |
| T1-T2 | 334 (48.1) | 164 (82.0) | 57 (62.6) | 30 (61.2) |  |
| T3-T4 | 361 (51.9) | 36 (18.0) | 34 (37.4) | 19 (38.8) |  |
| **cN (%)** |  |  |  |  | **<0.001** |
| N0 | 58 (8.3) | 35 (17.5) | 27 (29.7) | 16 (32.7) |  |
| N1-N3 | 637 (91.7) | 165 (82.5) | 64 (70.3) | 33 (67.3) |  |
| **HR status (%)** |  |  |  |  | 0.777 |
| Negative | 209 (30.1) | 59 (41.8) | 24 (26.4) | 17 (53.1) |  |
| Positive | 486 (69.9) | 141 (58.2) | 67 (73.6) | 32 (46.9) |  |
| **HER2 status (%)** |  |  |  |  | **<0.001** |
| Negative | 479 (68.9) | 179 (89.5) | 57 (62.6) | 27 (55.1) |  |
| Positive | 216 (31.1) | 21 (10.5) | 34 (37.4) | 22 (44.9) |  |
| **Subtype (%)** |  |  |  |  | **<0.001** |
| HR+/HER2- | 370 (53.2) | 134 (67.0) | 44 (48.4) | 15 (30.6) |  |
| HER2+ | 216 (31.1) | 20 (10.0) | 34 (37.4) | 22 (44.9) |  |
| TNBC | 109 (15.7) | 46 (23.0) | 13 (14.2) | 12 (24.5) |  |
| **Ki-67 index (%)** |  |  |  |  | **0.002** |
| Low (<20%) | 100 (14.4) | 12 (6.0) | 18 (19.8) | 7 (14.3) |  |
| High (≥20%) | 595 (85.6) | 188 (94.0) | 68 (74.7) | 42 (85.7) |  |
| Unknow | - | - | 5 (5.5) | - |  |
| **NG (%)** |  |  |  |  | **<0.001** |
| 1/2 | 486 (69.9) | 162 (81.0) | 79 (86.8) | 42 (85.7) |  |
| 3 | 209 (30.1) | 38 (19.0) | 12 (13.2) | 7 (14.3) |  |
| **sTILs (%)** |  |  |  |  | 0.195 |
| Low | 397 (57.1) | 125 (62.5) | 53 (58.2) | 20 (40.8) |  |
| Moderate | 251 (36.1) | 64 (32.0) | 30 (33.0) | 25 (51.0) |  |
| High | 47 (6.8) | 11 (5.5) | 8 (8.8) | 4 (8.2) |  |
| **pCR (%)** |  |  |  |  | **<0.001** |
| Yes | 169 (24.3) | 35 (17.5) | 37 (40.7) | 13 (26.5) |  |
| No | 526 (75.7) | 165 (82.5) | 54 (59.3) | 36 (73.5) |  |

*Abbreviations:* HR, hormone receptor; HER2, human epidermal growth factor receptor 2; sTILs, stromal tumor-infiltrating lymphocytes; NG, nuclear grade; pCR, pathological complete response.
Bold indicates statistical significance (P < 0.05)

## Results

### Clinical characteristics

Figure 1 shows the workflow of the patient recruitment. According to the inclusion and exclusion criteria, we enrolled a total of 1035 patients from four independent institutions: West China Hospital (WC cohort, 695 patients from 2010.04∼2021.04), Shanxi Cancer Hospital (SX cohort, 200 patients from 2015.02∼2019.10), Sichuan Province People’s Hospital (SC cohort, 91 patients from 2020.01∼2021.02), the Affiliated Hospital of Southwest Medical University (SW cohort, 49 patients from 2016.08∼2020.10). The dataset from West China Hospital had the largest population of eligible patients (N=695) and was used as the PC. The clinical characteristics of all patients are summarized in Table 1 (detail information was available in supplementary Table S3).

The pCR rate among the four cohort were between 17.5% to 40.7% (Table 1). As showing in Table S3, sTILs was significantly different between pCR and non-pCR group in all the four cohorts (*P* < 0.05). Besides, pCR was associated with HR status and subtype in all cohorts except V3. HER2 and NG were differently distributed between the two groups in PC and one validation cohort (V2, V1). However, pCR was significantly correlated with Ki67 and cT only in PC but not in the other three validation cohorts. We did not found significant difference of age and cN between pCR and non-pCR groups. Hence, Subtype, NG, Ki67, and cT were baseline predictors of pCR while sTILs was a strongly predictive factor manually evaluated from tumor-associated stroma.

### Automated detection of stromal compartment

The E-S classifier (CNN I) was applied to detect the stromal regions of all tiles cropped from ROIs of each WSI. A total of 55,078 tiles were generated from 1035 WSIs. Heat-map of stroma generated by CNN I was shown in Figure 2 and Figure S2. As a result, it achieved the accuracy of 0.806 and 0.827 for the stroma identification in the validation and testing set, respectively. Furthermore, the E-S classifier showed high precision of 0.896 and 0.870, which indicated that more that 85% area identified as stroma was exactly correct. After manually insepction, 44 stromal tiles per patient were enrolled on average (details were shown in Supplementary Materials, Figure S3, and Table S2). All remained tiles were delivered to the following experiments of developing a stroma-derived predictor.

### TS-score construction and validation

The construction pipeline of TS-score was depicted in Figure 2. A total of 44,734 stromal tiles with double certification from the E-S classifier and human observer were used. The Inception-V4 architecture was trained by learning from the stromal tiles with given labels of pCR or non-pCR in the primary cohort, and a 5-fold cross validation was used to determine the parameters of the CNN II. After scoring all tiles, the TS-score of a given patient was obtained from calculating the mean value of the tile-level, which reflected the predictive risk of getting a pCR based on the tumor stromal compartment. The ROC curves and AUCs of the raw TS-scores in the PC and three external validations are shown in Figure 3. The TS-score achieved an AUC of 0.729 to predict pCR in the PC and 0.745, 0.673, 0725 in the V1, V2, V3 datasets at WSI-level, respectively. Additionally, TS-score showed stable performance on HR+HER2-breast cancer (AUC of subtype 1: PC 0.767, VC1 0.804, VC2 0.784, VC3 1.00), while patch-level performance of TS-score according to the three breast cancer subtypes are also shown in Figure 3. Detail results were available in Table S4 and S5.

**Figure 3.**
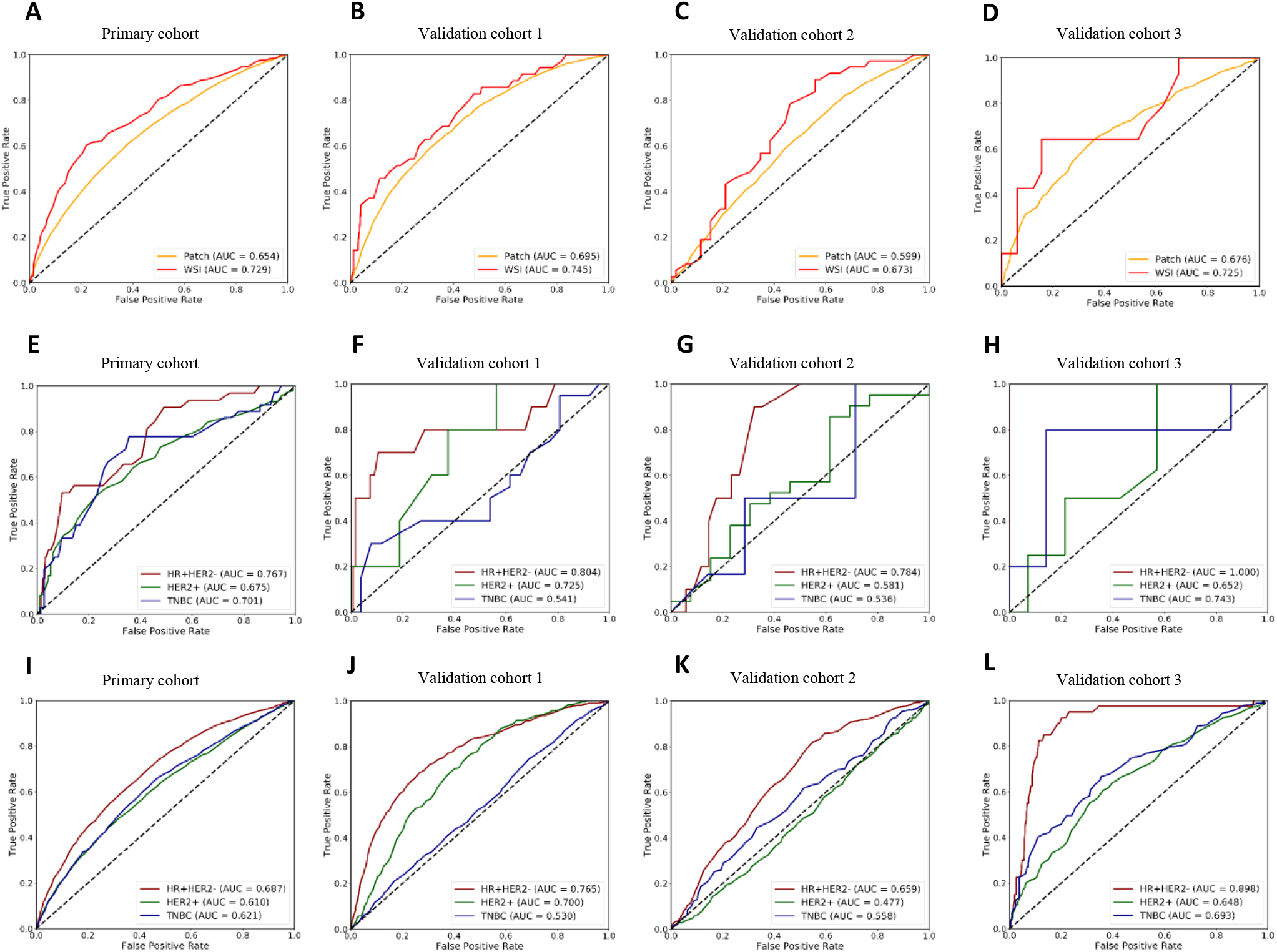
ROC curves of TS-score in the four hospitals. A, B, C, D show the WSI-level and patch-level performance of TS-score in the total dataset among the four centers; E, F, G, H show the WSI-level performance of TS-score in different breast cancer subtypes, and I, J, K, L show the patch-level performance of TS-score in different breast cancer subtypes.

### TS-score is independent of clinical variables and improves the prediction of pCR

To evaluate the independent predictive power of pCR risk by TS-score, we performed the multivariate logistic regression analysis including factors which were potentially correlated with pCR (Table 2 and Table S6) in the four datasets; due to the limited data size of the external validations, we combined the three validation cohorts to perform the following analysis. As results of PC showing in Table 2, TS-score was significantly correlated with pCR in univariate analysis (*P* < 0.001), and it remained predictive when correcting for all other factors, including sTILs, subtype, T stage, Ki67, and nuclear grade (*P* < 0.001). Subtype was also significant (*P* < 0.001) but sTILs was not (*P* = 0.766), although sTILs was indeed a significant predictor in multivariate analysis without TS-score (*P* < 0.001). The similar results were observed in the external validations, as that the TS-score was an independent predictor of pCR (*P* = 0.013) (Table S6). Furthermore, using the logistic regression method we developed factor-based prediction models of pCR to compare the predictive ability of TS-score with other CP factors (Figure 4A and 4B). TS-score based model yielded the best performance with an AUC of 0.727 in the PC, which was comparable to subtype-based model (AUC = 0.727, *P* = 0.927) and even outperformed the sTILs-based model (AUC = 0.651, *P* < 0.001), similarly in the validations. Detailed results were available in Table S7.

**Table 2.** Univariate and multivariate analysis of TS-score correlating with pCR in primary cohort.

| Factors | Univariate analysis |  | Multivariate analysis <sup>1</sup> |  | Multivariate analysis <sup>2</sup> |  |
| --- | --- | --- | --- | --- | --- | --- |
|  | OR(95% CI) | P | OR(95% CI) | P | OR(95% CI) | P |
| TS-score | - | **<0.001** | - | - | - | **<0.001** |
| Subtypes | - | **<0.001** | - | **<0.001** | - | **<0.001** |
| HR+/HER2- | 1 | - | 1 | - | 1 | - |
| HER2+ | 9.28(5.91-14.6) | **<0.001** | 7.47(4.66-12.0) | **<0.001** | 7.73(4.76-12.5) | **<0.001** |
| TNBC | 5.21(3.04-8.93) | **<0.001** | 3.73(2.10-6.62) | **<0.001** | 3.33(1.86-5.97) | **<0.001** |
| sTILs | - | **<0.001** | - | **<0.001** | - | 0.766 |
| Low | 1 | - | 1 | - | 1 | - |
| Moderate | 7.58(4.00-14.4) | **<0.001** | 1.76(1.16-2.69) | **0.009** | 1.03(0.64-1.66) | 0.905 |
| High | 2.78(1.47-5.25) | **0.002** | 4.58(2.20-9.54) | **<0.001** | 1.36(0.57-3.27) | 0.490 |
| cT | 0.61(0.43-0.86) | **0.005** | 0.73(0.49-1.10) | 0.130 | 0.77(0.51-1.16) | 0.204 |
| Ki67 | 2.63(1.40-4.94) | **0.003** | 1.40(0.69-2.81) | 0.348 | 1.19(0.58-2.46) | 0.636 |
| NG | 2.59(1.80-3.70) | **<0.001** | 1.37(0.91-2.07) | 0.137 | 1.21(0.79-1.85) | 0.372 |

*Note:* Multivariate analysis <sup>1</sup> refers to the multivariate analysis excluding TS-score; Multivariate analysis <sup>2</sup> refers to the multivariate analysis including the TS-score.
*Abbreviation:* TS-score, tumor-stroma score; HR, hormone receptor; HER2, human epidermal growth factor receptor 2; sTILs, stromal tumor-infiltrating lymphocytes; NG, nuclear grade
Bold indicates statistical significance ( $P < 0.05$ )

**Figure 4.**
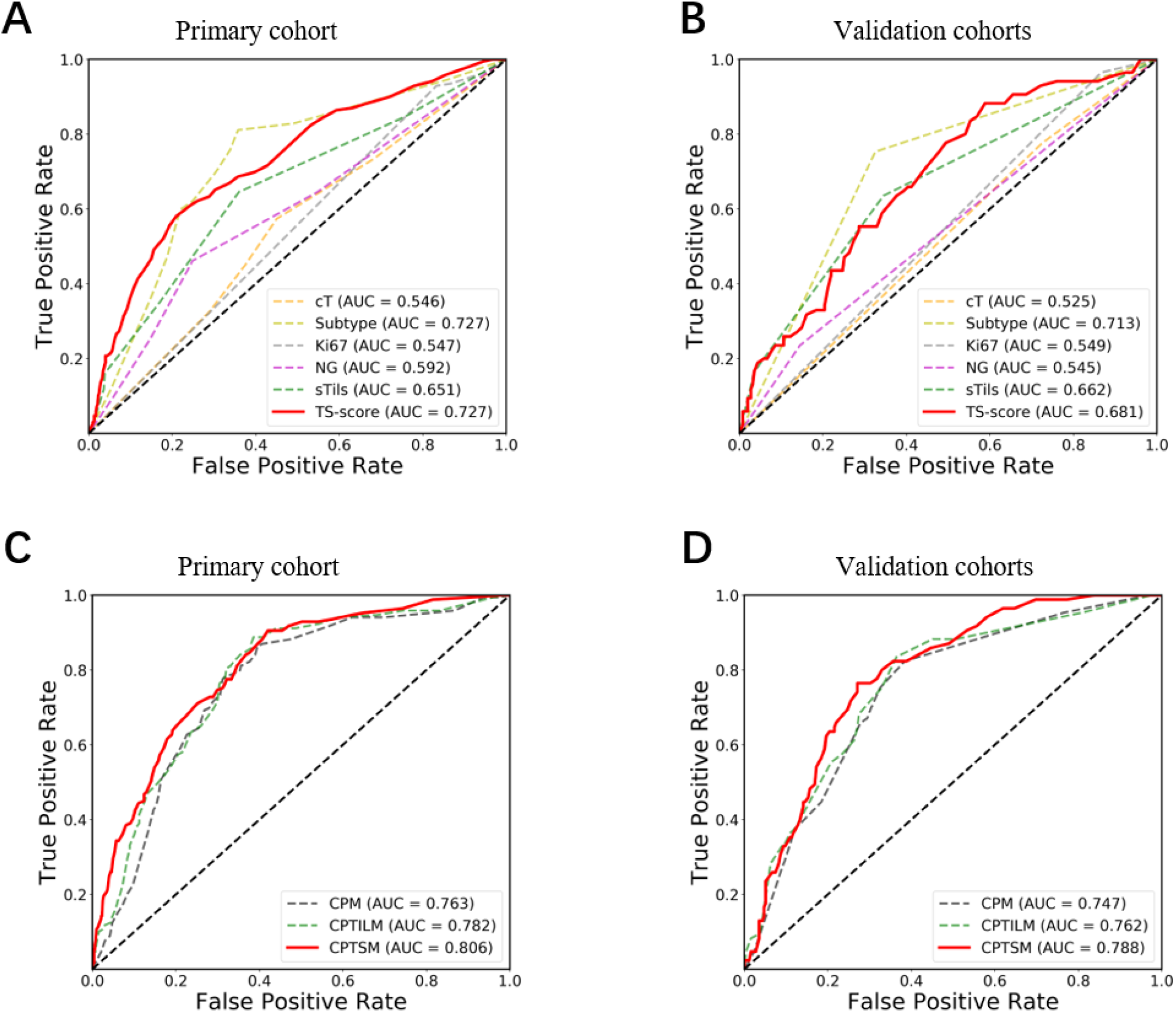
ROC curves of the marker-based models. The top row shows the performance of the single marker-based models for predicting pCR in the primary cohort (A) and the three external validations (B). The bottom row shows the performance of the baseline marker-based model (CPM), the baseline marker and sTILs-based model (CPTILM), and the baseline marker and TS-score-based model (CPTSM) for predicting pCR in three three external validations (C, D).

**Figure 5.**
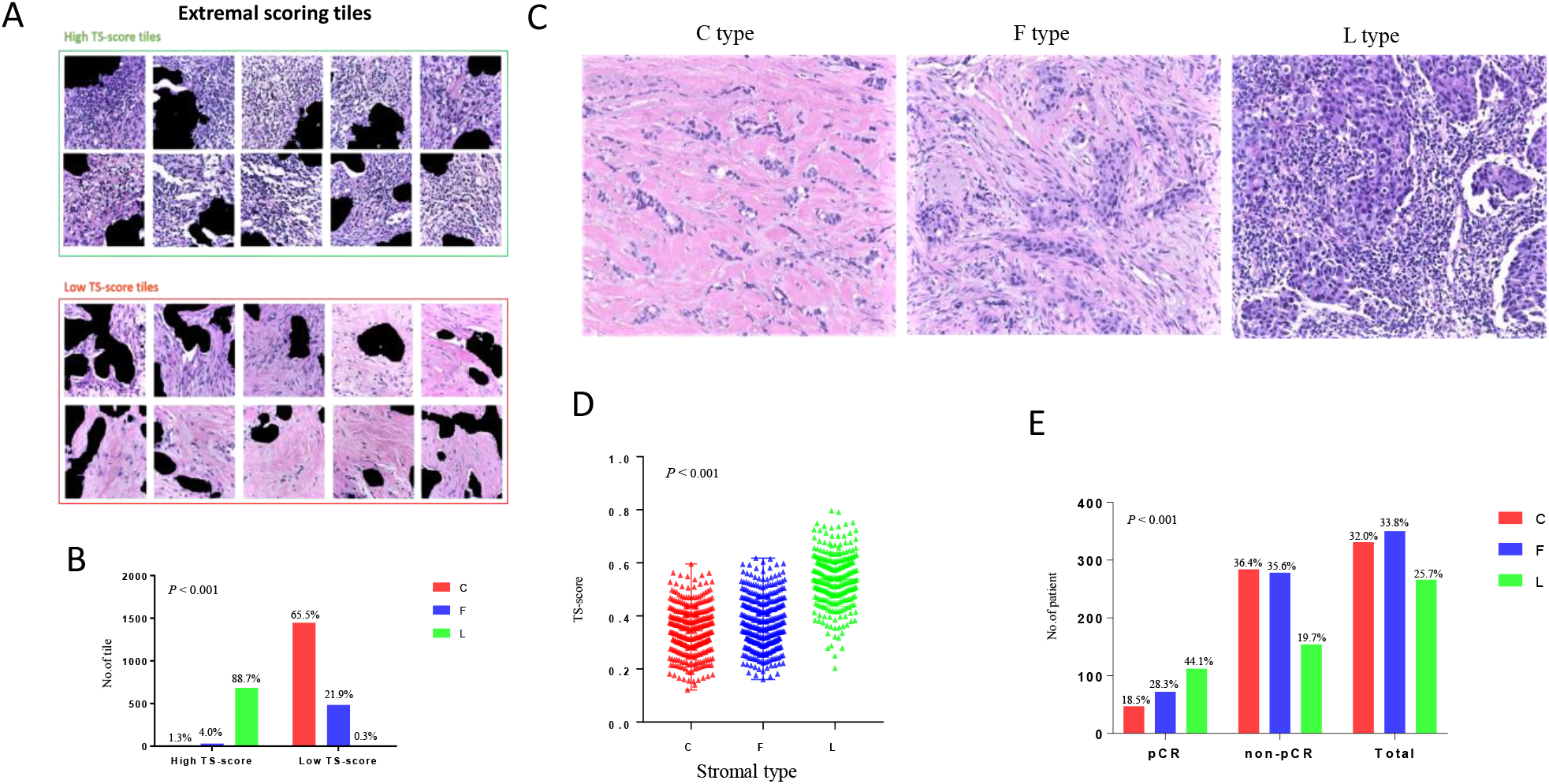
The underlying histological patterns of TS-score characterizing at the patch-level (A, B) and WSI-level (C, D, E). A. Tiles with extremal TS-scores associated with pCR and non-pCR were extracted to be reviewed by a pathologist. Scale bars, 233μm. B. The distribution of tiles with different stromal type between extremal high TS-score and low TS-score group. C. Examples of different stromal type: collagen-dominant stromal (C type), fibroblast-dominant stroma (F type), lymphocyte-dominant stroma (L type). Scale bars, 100μm. D. The distributions of TS-scores among the three stromal type evaluated at WSI-level. E. The different percentage of the three stromal type between pCR and non-pCR group.

Univariate logistic regression analysis revealed that baseline CP factors, including subtype, NG, Ki67, and cT, were significantly predictive of pCR; sTILs and TS-score, stroma-derived factors, were predictive as well. Therefore, a CP-based model combining subtype, NG, Ki67, and cT was constructed using the primary cohort (CPM); meanwhile, a model combined these factors above and TS-score was developed to evaluate the prediction incremental value of TS-score (CPTSM). Additionally, a prediction model combining CP factors and sTILs was built as a comparison (CPTILM). As shown in the Figure 4C, CPTSM yielded an AUC of 0.806 in the PC, while an AUC of 0.763 for CPM and 0.782 for CPTILM. Using the DeLong test, the CPTSM showed significantly higher AUC than that of both the CPM (*P* < 0.001) and CPTILM (*P* = 0.005) (Table 3). Similar results were also observed in the external validations; the CPTSM significantly outperformed the CPM (*P* =0.027), and also showed higher AUC than CPTILM with close to a significance (*P =* 0.078) (Figure 4D, Table 3). Results for VC1, VC2 and VC3 were available in Figure S6 and Table S8.

**Table 3.** AUCs of models for pCR prediction in the primary and validation cohorts

|  | CPTSM (95% CI) | CPTILM (95% CI) | CPM (95% CI) |
| --- | --- | --- | --- |
| PC | 0.806 (0.78-0.83) | 0.782 (0.73-0.83) | 0.763 (0.72-0.81) |
| p value<sup>1</sup> | **<0.001** | **0.04** | 1 |
| p value<sup>2</sup> | **0.005** | 1 | - |
| VCs | 0.788 (0.783-0.793) | 0.762 (0.756-0.768) | 0.747 (0.742-0.752) |
| p value<sup>1</sup> | **0.027** | 0.180 | 1 |
| p value<sup>2</sup> | 0.078 | 1 | - |

*Abbreviations:* AUC, area under receiver operating characteristic curve; pCR, pathological complete response; CPM, clinicopathology-based model; CPTILM, clinicopathology and sTILs based model; CPTSM, clinicopathology and TS-score based model.
p value refers to Delong test for the differences of AUCs between different metrics in different cohorts; p value<sup>1</sup> refers the comparisons with the CPM while p value<sup>2</sup> refers the comparisons with CPTILM.
Bold indicates statistical significance ( $P < 0.05$ )

### TS-score reflects the stromal histological patterns which correlated with pCR

To obtain an overall understanding of the histological patterns which contributes to the exact prediction, the distributions of each tile score in the PC were visualized and the tiles corresponding to extremal scores (top 10% and the bottom 10%) were extracted for manual evaluation. These extremal patches (n=2,980) were classified into three stromal phenotypes, which were C type, F type, and L type [27]; tiles which did not belong to the three type were excluded of analysis (Figure 5C). High TS-score tiles were mainly L type while stroma of C type was few (684/771, 10/771). In contrast, low TS score tiles mainly showed rich collagen or partly had a higher distribution of F type-stroma while with an extremely low percentage of L type-stroma (1447/2,209, 484/2,209,7/2,209). Significant difference was showed among the distributions of stromal type between tiles of the highest 10% and lowest 10% TS-score (P < 0.001) (Figure 5A,5B), as high scores are mostly predicted on lymphocyte dominant regions and low scores are mainly for collagen dominant stroma.

To further determine the relationship between stromal histological patterns and treatment response to NAC, we also assessed the stromal type of 1035 HE-stained images among the four hospitals at WSI-level. In pCR group, L type-stroma was the dominant (44.1%) while patients of C type and F-type stroma were more in non-pCR group (36.4%, 35.6%, P < 0.001) (Figure 5E). Correspondingly, cases of L type-stroma showed the highest TS-score, F type is next, and C type is the lowest (P < 0.001) (Figure 5D). Additionally, the pCR rate was 14.2%, 20.6%, and 42.1% in cases of C type-stroma, F type-stroma, and L type-stroma, respectively (Table S9).

## Discussion

In this study, we proposed a new stromal derived biomarker, TS-score, and investigated its predictive ability for treatment response to NAC with a multi-center dataset. Experiments showed that TS-score is predictive of pCR independently of subtype, tumor size, sTILs, nuclear grade, and Ki67, which can provide complementary information for predicting pCR outperforming the routine clinic-pathological biomarkers. According to the histological patterns reflected by TS-score, interestingly, we discovered that stroma with collagen and fibroblast dominant was likely associated with inadequate response to NAC, which was contrary to lymphocyte dominant stroma. We also assessed the stromal type on WSI and further identified this relationship between stromal histological patterns and pCR at the patient-level. In summary, TS-score, which is directly obtained from routine HE-stained images, can serve as a potential candidate that improves the prediction of pCR in breast cancer.

Our study investigated the DL-based prediction of pathological complete response to NAC for breast cancer, a disease with the highest incidence in female and wide variations in treatment response to NAC [38]. Treatment planning for breast cancer is dependent on several factors, such as clinical stage and molecular subtype. Due to their limited prediction, the field of breast oncology is currently awaiting features that can better distinguish chemotherapy-sensitivity and chemotherapy-resistance. In our study, TS-score outperformed the baseline predictors in predicting pCR and performed as well as molecular subtype in the primary and external validation cohorts, despite using only a very small portion of each histological images. Remarkably, TS-score even outperformed sTILs despite the fact that both are stromal histological predictors for pCR. Given that histological assessment of sTILs has been limited by considerable intra- and inter-observer variability, TS-score can effectively extract the predictive information from histological images via a highly reproducible and quantitative approach that compensates for the defect of sTILs. Additional investigations of the independence of TS-score revealed that it can provide complementary information to the baseline factors for predicting pCR and the comparisons of models demonstrated that the addition of TS-score can improve the prediction performance with statistic significance, which is meaningful as that improving the prediction can facilitate the favorable patient care in NAC settings.

In breast cancer, the potential ability of tumor-associated stroma was investigated via manual pathological evaluation [22-24, 29], molecular biological assays [39, 40], and digital pathological techniques [18, 41], showing that stroma could facilitate for disease classification and outcome prediction. However, to our knowledge, the potential information in the stromal compartment has not been mined to predict the pCR by DL techniques. Therefore, our study constitutes a precedent for objectively assessing the hidden information from the stroma and proposing a stroma-derived biomarker to improve the prediction of pCR in breast cancer. Conventionally, pathological diagnosis is based largely on the histological appearance and molecular characteristics of epithelial cells, while stroma alterations are often subtle and difficult to characterize and quantify by manual evaluation. Moreover, tumor stroma is highly heterogeneous and complex in breast cancer, which could be challengeable for the ROIs selection and automated identification of tumor stroma. Hence, manual annotation was required in our study to select the representative regions like a previous study in prostate cancer [19]. Unlike the the automated detection for tumor epithelium [20], stroma identification is a more difficult task; hence, we employed the state of art algorithms and used a larger sample size to construct an improved model to detect the stromal pixels within ROIs. Unlike Kather and colleagues [17], who quantified the various components of the stroma in colorectal cancer and combined them into a stroma score, the CNN II in our study learned directly from the stromal compartment, integrating the predictive information into a biomarker: the TS-score. Compared to their study, an end-to-end approach to extract information is likely simpler and more potentially to discover the hidden interaction patterns between different components despite the weakened interpretability.

Another important aspect of TS-score is its interpretability. As in all studies using DL-based methods, one question always arises: what the output score represents exactly. Essentially, many AI-based models or biomarkers are black boxes, which are difficult to explain their predictions in the way that human can understand. This is crucial, however, as these will be widely used and supported only when the underlying decision process can be understood. The tile scoring procedure allows us to score each tile of a given WSI, guiding pathologists to stromal patterns associated with treatment response. The extremal tiles suggest that TS-score can reflect the known histological features related to pCR, such as sTILs. The abundance of sTILs is associated with treatment response to NAC in breast cancers, and patients with high density of sTILs are more likely to achieve a pCR [22, 23]. Correspondingly, tiles of lymphocyte dominant stroma were assigned with a high TS-score, which indicates that the CNN network perceives the feature of sTILs abundance in the stroma. On the other hand, tiles of collagen and fibroblast dominant stroma were predicted with low TS-scores, indicating a high risk of missing pCR, but these histological patterns have not been generally recognized as features of chemo-resistance behavior or incorporated into pathological evaluating paradigms. These findings guided us to further clarify the relationship between stromal histological patterns and treatment response through manual classification. Although the dominant stromal type has shown to be predictive for prognosis in breast cancer [27], we provided the evidence for its potential relationship with pCR.

In fact, AI technique-based image analysis has broad applications in the modern medicine. In radiology, some DL-based inventions have already been approved by FDA [42, 43]. Compared to these imaging modalities, histological images contain more abundant information and provide the gold standard for diagnosis; combining the AI techniques can have promising prospects for clinical use. Clinical translation of digital pathology, however, is still in its infancy. To advance clinical applications, large amounts of training data and robust multi-center validation are required, while many current studies are exactly hampered by these limitations. In the present study, we addressed the troubles: four independent datasets of more than one thousand cases were used to establish and validate the CNN-based TS-score as a predictive biomarker in breast cancer. With this approach, we showed the DL-based stromal score improved the prediction of pCR in breast cancer. Furthermore, validating it in three external datasets, we confirmed the predictive potentials of this approach. Therefore, we presented a novel candidate for NAC response prediction, which could be combined with existing predictors to promote the better stratification of patients and facilitate the clinical decision-making.

The study did have some limitations. Although 1035 breast cancer patients were collected from four hospitals, the size of the validation dataset was limited, with two validation cohorts including patients less than 100. Furthermore, we only included the retrospective data, while this study needs to be validated prospectively.

In spite of these limitations, our study is the first to show the potential ability of breast cancer stromal compartment in pCR prediction via a DL-based approach. Furthermore, the findings of this study provide some insight into different characteristics of TME between pCR and non-pCR breast cancer patients. Future work will need to replicate and validate these findings in larger cohorts and prospective clinical trials. In addition, we will continue our studies on the spatial patterns between tumor epithelium and stroma to further explore the potentials of the breast cancer histology.

## Data and code availability

Data and related codes used and/or analyzed during the current study are available from the corresponding author on reasonable request.

## Acknowledgement

This work was supported by the 1·3·5 project for disciplines of excellence (ZYGD18012); the Technological Innovation Project of Chengdu New Industrial Technology Research Institute (2017-CY02–00026-GX).

## Author Contributions

H.B., Z.Z., Y.Y., and F.L. designed and supervised this project. F.L., Y.W., Y.Z., J.F., and X.X. collected the data used in this study. Y.Y. and F.L. completed the data analysis and interpretation. F.L. wrote the initial paper. Y.Y., Z.Z., and H.B. edited the paper. All authors discussed the results and approved the paper.

## References

1. Derks MGM, van de Velde CJH. Neoadjuvant chemotherapy in breast cancer: more than just downsizing. Lancet Oncol 2018; 19: 2–3.

2. von Minckwitz G, Blohmer JU, Costa SD et al. Response-guided neoadjuvant chemotherapy for breast cancer. J Clin Oncol 2013; 31: 3623–3630.

3. Cortazar P, Zhang L, Untch M et al. Pathological complete response and long-term clinical benefit in breast cancer: the CTNeoBC pooled analysis. Lancet 2014; 384: 164–172.

4. Cain H, Macpherson IR, Beresford M et al. Neoadjuvant Therapy in Early Breast Cancer: Treatment Considerations and Common Debates in Practice. Clin Oncol (R Coll Radiol) 2017; 29: 642–652.

5. Haque W, Verma V, Hatch S et al. Response rates and pathologic complete response by breast cancer molecular subtype following neoadjuvant chemotherapy. Breast Cancer Res Treat 2018; 170: 559–567.

6. Echle A, Rindtorff NT, Brinker TJ et al. Deep learning in cancer pathology: a new generation of clinical biomarkers. Br J Cancer 2020; 124: 686–696.

7. Acs B, Rantalainen M, Hartman J. Artificial intelligence as the next step towards precision pathology. J Intern Med 2020; 288: 62–81.

8. Simonyan K, Zisserman A. Very deep convolutional networks for large-scale image recognition. arXiv preprint 1409.1556 2014.

9. Ehteshami Bejnordi B, Veta M, Johannes van Diest P et al. Diagnostic Assessment of Deep Learning Algorithms for Detection of Lymph Node Metastases in Women With Breast Cancer. Jama 2017; 318: 2199–2210.

10. Bulten W, Pinckaers H, van Boven H et al. Automated deep-learning system for Gleason grading of prostate cancer using biopsies: a diagnostic study. Lancet Oncol 2020; 21: 233–241.

11. Campanella G, Hanna MG, Geneslaw L et al. Clinical-grade computational pathology using weakly supervised deep learning on whole slide images. Nat Med 2019; 25: 1301–1309.

12. Akbar S, Jordan LB, Purdie CA et al. Comparing computer-generated and pathologist-generated tumour segmentations for immunohistochemical scoring of breast tissue microarrays. Br J Cancer 2015; 113: 1075–1080.

13. Coudray N, Ocampo PS, Sakellaropoulos T et al. Classification and mutation prediction from non-small cell lung cancer histopathology images using deep learning. Nat Med 2018; 24: 1559–1567.

14. Courtiol P, Maussion C, Moarii M et al. Deep learning-based classification of mesothelioma improves prediction of patient outcome. Nat Med 2019; 25: 1519–1525.

15. Woerl AC, Eckstein M, Geiger J et al. Deep Learning Predicts Molecular Subtype of Muscle-invasive Bladder Cancer from Conventional Histopathological Slides. Eur Urol 2020; 78: 256–264.

16. Zhang F, Yao S, Li Z et al. Predicting treatment response to neoadjuvant chemoradiotherapy in local advanced rectal cancer by biopsy digital pathology image features. Clin Transl Med 2020; 10.

17. Kather JN, Krisam J, Charoentong P et al. Predicting survival from colorectal cancer histology slides using deep learning: A retrospective multicenter study. PLoS Med 2019; 16: e1002730.

18. Beck AH, Sangoi AR, Leung S et al. Systematic analysis of breast cancer morphology uncovers stromal features associated with survival. Sci Transl Med 2011; 3: 108ra113.

19. Bhargava HK, Leo P, Elliott R et al. Computationally Derived Image Signature of Stromal Morphology Is Prognostic of Prostate Cancer Recurrence Following Prostatectomy in African American Patients. Clin Cancer Res 2020; 26: 1915–1923.

20. Li F, Yang Y, Wei Y et al. Deep learning-based predictive biomarker of pathological complete response to neoadjuvant chemotherapy from histological images in breast cancer. J Transl Med 2021; 19: 348.

21. Conklin MW, Keely PJ. Why the stroma matters in breast cancer: insights into breast cancer patient outcomes through the examination of stromal biomarkers. Cell Adh Migr 2012; 6: 249–260.

22. Denkert C, von Minckwitz G, Darb-Esfahani S et al. Tumour-infiltrating lymphocytes and prognosis in different subtypes of breast cancer: a pooled analysis of 3771 patients treated with neoadjuvant therapy. Lancet Oncol 2018; 19: 40–50.

23. Denkert C, Loibl S, Noske A et al. Tumor-associated lymphocytes as an independent predictor of response to neoadjuvant chemotherapy in breast cancer. J Clin Oncol 2010; 28: 105–113.

24. Hagenaars SC, de Groot S, Cohen D et al. Tumor-stroma ratio is associated with Miller-Payne score and pathological response to neoadjuvant chemotherapy in HER2-negative early breast cancer. Int J Cancer 2021; 149: 1181–1188.

25. Goldhirsch A, Winer EP, Coates A et al. Personalizing the treatment of women with early breast cancer: highlights of the St Gallen International Expert Consensus on the Primary Therapy of Early Breast Cancer 2013. Annals of oncology 2013; 24: 2206–2223.

26. Salgado R, Denkert C, Demaria S et al. The evaluation of tumor-infiltrating lymphocytes (TILs) in breast cancer: recommendations by an International TILs Working Group 2014. Ann Oncol 2015; 26: 259–271.

27. Ahn S, Cho J, Sung J et al. The prognostic significance of tumor-associated stroma in invasive breast carcinoma. Tumour Biol 2012; 33: 1573–1580.

28. de Kruijf EM, van Nes JG, van de Velde CJ et al. Tumor-stroma ratio in the primary tumor is a prognostic factor in early breast cancer patients, especially in triple-negative carcinoma patients. Breast Cancer Res Treat 2011; 125: 687–696.

29. Dekker TJ, Charehbili A, Smit VT et al. Disorganised stroma determined on pre-treatment breast cancer biopsies is associated with poor response to neoadjuvant chemotherapy: Results from the NEOZOTAC trial. Mol Oncol 2015; 9: 1120–1128.

30. Yang Y. One-Step Abductive Multi-Target Learning with Diverse Noisy Samples. arXiv preprint 2110.10325 2021.

31. Szegedy C, Ioffe S, Vanhoucke V, Alemi AA. Inception-v4, inception-resnet and the impact of residual connections on learning. In Thirty-first AAAI conference on artificial intelligence. 2017.

32. Aurelio YS, de Almeida GM, de Castro CL, Braga AP. Learning from imbalanced data sets with weighted cross-entropy function. Neural processing letters 2019; 50: 1937–1949.

33. Theodoridis S. Chapter 5 - Stochastic Gradient Descent: The LMS Algorithm and its Family. In Theodoridis S (ed) Machine Learning. Oxford: Academic Press 2015; 161–231.

34. Yang Y, Lv H, Chen N et al. Local Minima Found in the Subparameter Space Can Be Effective for Ensembles of Deep Convolutional Neural Networks. Pattern Recognition 2020; 109: 107582.

35. Yongquan Y, Haijun L, Ning C et al. FTBME: feature transferring based multi-model ensemble. Multimedia Tools and Applications 2020; 79: 18767–18799.

36. Yang Y, Lv H, Chen N. A Survey on Ensemble Learning under the Era of Deep Learning. arXiv preprint 2101.08387 2021.

37. DeLong ER, DeLong DM, Clarke-Pearson DL. Comparing the areas under two or more correlated receiver operating characteristic curves: a nonparametric approach. Biometrics 1988; 837–845.

38. Siegel RL, Miller KD, Fuchs HE, Jemal A. Cancer statistics, 2022. CA: a cancer journal for clinicians 2022.

39. Farmer P, Bonnefoi H, Anderle P et al. A stroma-related gene signature predicts resistance to neoadjuvant chemotherapy in breast cancer. Nat Med 2009; 15: 68–74.

40. Finak G, Bertos N, Pepin F et al. Stromal gene expression predicts clinical outcome in breast cancer. Nat Med 2008; 14: 518–527.

41. Ehteshami Bejnordi B, Mullooly M, Pfeiffer RM et al. Using deep convolutional neural networks to identify and classify tumor-associated stroma in diagnostic breast biopsies. Mod Pathol 2018; 31: 1502–1512.

42. Ardila D, Kiraly AP, Bharadwaj S et al. End-to-end lung cancer screening with three-dimensional deep learning on low-dose chest computed tomography. Nat Med 2019; 25: 954–961.

43. Luo H, Xu G, Li C et al. Real-time artificial intelligence for detection of upper gastrointestinal cancer by endoscopy: a multicentre, case-control, diagnostic study. Lancet Oncol 2019; 20: 1645–1654.

